# OddCAPS: a simple, low-cost, universal technique for detecting single nucleotide variants

**DOI:** 10.64898/2026.07.16.738826

**Authors:** Karin Kawaguchi, Yuzuha Komachiya, Mai Muto, Ryoka Teshima, Naoko Sakai, Hayao Ohno

**Author notes:** Correspondence to (H.O.).

## Abstract

Derived Cleaved Amplified Polymorphic Sequences (dCAPS) assays have been widely performed historically to detect known base substitutions in many model organisms—notably *Caenorhabditis elegans*, *Arabidopsis thaliana*, and yeast—where chemical mutagens that induce point mutations are frequently used. With the rise of whole-genome sequencing and genome editing technologies, dCAPS is increasingly applied to detect diverse nucleotide changes across a broader range of species. However, a key limitation of dCAPS is that genomic target sites amenable to primer designs that both preserve PCR amplification and create recognition sites for inexpensive, high-performance restriction enzymes are scarce. Here we report One-step dual-primer dCAPS (OddCAPS), a modification that uses three primers in one reaction to overcome this constraint. Two of these primers, an intermediate primer and a dCAPS primer, sequentially introduce 1–2 base substitutions each into the amplicon, enabling up to four engineered base changes near the nucleotide of interest. By using the intermediate primer at 1/10–1/100 the concentration of the other primers, the desired product is generated directly in a single-tube, one-step PCR. Increasing the number of engineered substitutions improves the chance of using a researcher’s preferred restriction enzyme. In principle, having eight common restriction enzymes (BamHI, EcoRI, NheI, SalI, BglII, ClaI, HindIII, and MluI) suffices to detect any single-nucleotide variant in any biological or synthetic DNA sequence with this approach.

## Introduction

Detection of DNA base substitutions, including known point mutations and single nucleotide polymorphisms (SNPs), is an essential component of genotyping in molecular biology and genetics and is often applied to hundreds of samples. The emergence of techniques such as whole-genome sequencing and CRISPR-Cas9 has repeatedly generated a need for faster, less expensive methods to detect single nucleotide variants in DNA, and a variety of approaches have been developed to meet this need. (1) Allele-specific PCR (Newton et al., 1989; Touroutine & Tanis, 2020) employs primers designed to discriminate the nucleotide at the target position and determines the presence or absence of a substitution by the occurrence of PCR amplification. Because the assay relies solely on whether amplification occurs, it is rapid; however, achieving acceptable specificity often requires careful optimization of reaction conditions, and in many cases two separate PCR reactions and agarose gel electrophoresis lanes are needed—one that detects the presence of the variant and one that detects its absence. In addition, PCR is relatively prone to stochastic failure compared with other enzymatic assays (for example restriction digests), so methods that call genotype solely on amplification success risk incorrect calls if wells fail to amplify for non-biological reasons (e.g., partial evaporation). (2) Probe-based real-time PCR assays such as TaqMan (Livak, 1999) offer high specificity and quantitative capability, but they require design and synthesis of a fluorescently labeled probe for each target, which increases cost. (3) High-throughput methods such as kompetitive allele specific PCR (KASP) (Semagn et al., 2013) and high-resolution melting (HRM) analysis (Wittwer et al., 2003) scale well but demand specialized fluorescence detection instruments and often expensive proprietary reagents. (4) PCR-restriction fragment length polymorphism (PCR-RFLP) or cleaved amplified polymorphic sequences (CAPS) assays (Konieczny & Ausubel, 1993) are simple and low-cost but can only be applied when a variant of interest creates or abolishes a restriction enzyme recognition site.

As an extension of PCR-RFLP, the dCAPS method (Neff et al., 1998) introduces intentional mismatches in primers to create one or two nucleotide changes in the PCR product that generate a restriction site. Its robustness, broad applicability, procedural simplicity, and the fact that it can be performed using only a conventional thermal cycler and agarose gel electrophoresis system have made dCAPS widely adopted. Nevertheless, in many cases primers cannot be designed successfully for particular target sequences or the available restriction enzymes are not commonly used. For example, two previous studies (Ghanizadeh et al., 2021; Neff et al., 1998) that used dCAPS employed the enzymes MnlI, BslI, HinfI, XcmI, BstAPI, AflII, MaeIII, MscI, Sau96I, and NlaIV. These are not the typical, broadly stocked restriction enzymes and may require new purchases; they are frequently much more expensive than common enzymes such as EcoRI. For purchases from New England Biolabs (NEB), the price per unit for these enzymes ranges from 2.5– to 92-fold that of EcoRI (https://www.neb.com; MaeIII is unavailable from NEB). This reliance on relatively expensive enzymes increases the cost per sample, a particular burden for large-scale genotyping.

We report an extension of the dCAPS approach in which an intermediate primer, supplied at 1/10–1/100 of the normal primer concentration, enables amplification of PCR products bearing up to four engineered base substitutions. The protocol allows genotyping (wild type versus mutant versus heterozygote) from a single PCR and a single agarose gel lane and increases the chance of generating restriction sites recognizable by inexpensive, routinely stocked restriction enzymes— thereby lowering consumable costs. Given the substantial decline in oligonucleotide synthesis costs relative to enzyme prices, this approach achieves both robustness and economy. In this study we validate the approach by genotyping mutations in the nematode *Caenorhabditis elegans*. Theoretically, however, any single-nucleotide variant in any biological or synthetic DNA sequence should be detectable using only eight widely available restriction enzymes (BamHI, EcoRI, NheI, SalI, BglII, ClaI, HindIII, and MluI).

## Results

As an example, consider detecting the presence of the terminal cytosine in the genomic DNA sequence 5’-ACACTATCTGGATGGTCAAAAAGTGGAGCAGATGCT***C***-3’ (the target base C is shown in italics). In the conventional dCAPS approach one would perform PCR using a primer that introduces a mismatch near the 3’ end, for instance 5’-ACACTATCTGGATGGTCAAAAAGTGGAGCAGATG<u>A</u>T***C***-3’ (the mismatch base is underlined). The resulting PCR product contains the 5’-G<u>A</u>T***C***-3’ sequence and can therefore be digested with the restriction enzyme MboI (Fig. 1A); successful cleavage serves as an indicator of the target C. However, MboI is relatively costly—about 16 times the price per unit of EcoRI when purchased from NEB (https://www.neb.com/)—and, as a 4-bp cutter, it is prone to cleaving at additional nearby sites, which can compromise detection of the base substitution. Moreover, MboI is reported to lose activity during prolonged incubations (https://www.neb.com/en/tools-and-resources/usage-guidelines/restriction-endonucleases-survival-in-a-reaction), and its instability can be problematic in workflows that require adding the restriction enzyme directly to the PCR mix for convenience, since insufficient digestion may result under such conditions.

**Figure 1.**
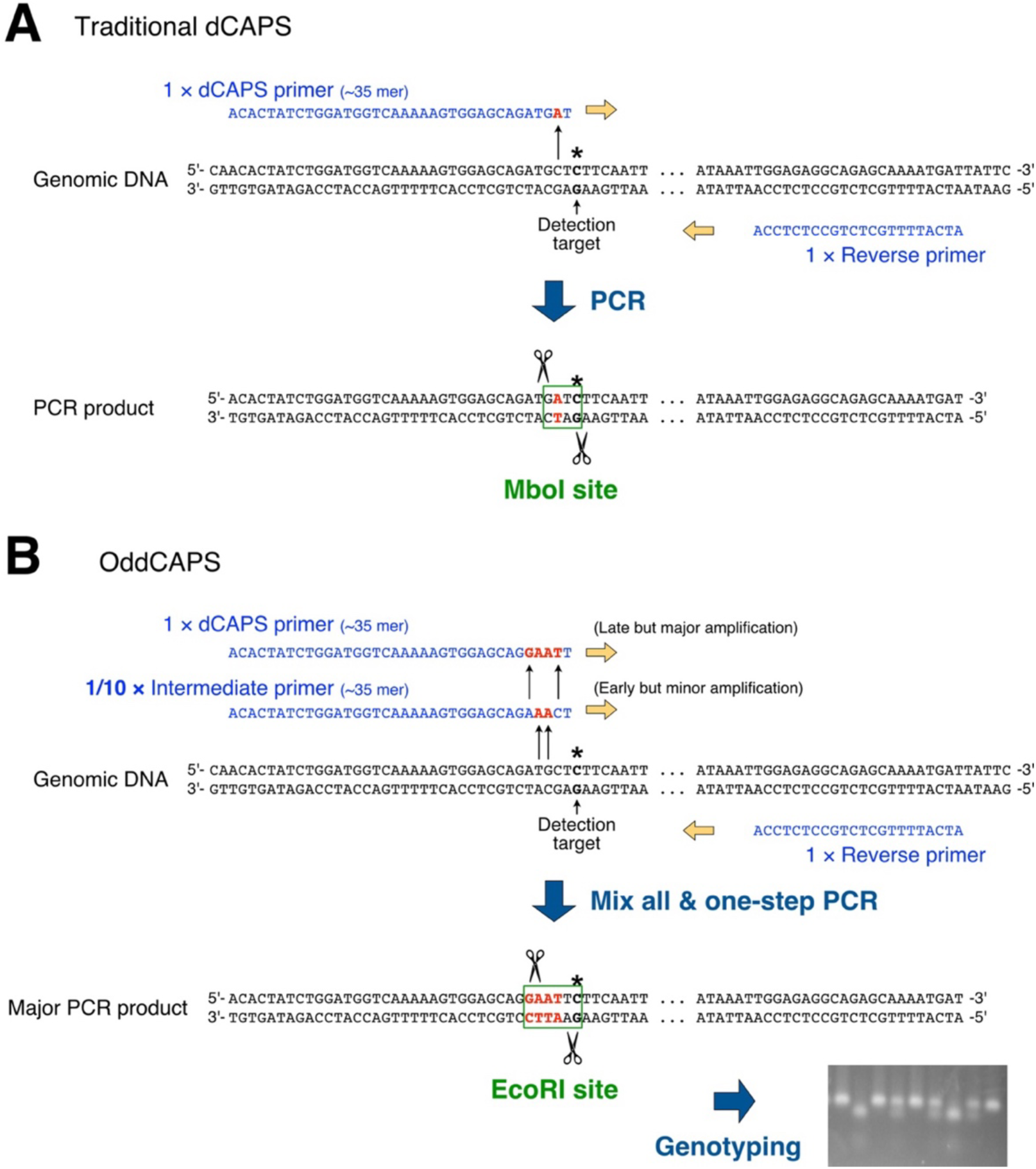
Schematic of detecting a specific nucleotide in genomic DNA using the OddCAPS method. (**A**) Example of detection by the dCAPS method. PCR is performed with a dCAPS primer and a reverse primer. (**B**) Example of detection by the OddCAPS method. PCR is performed with a mixture of a dCAPS primer, a diluted intermediate primer, and a reverse primer. (**A**, **B**) Red characters indicate base substitutions introduced by mismatches between the primers and the genomic template. The target base is indicated in bold with an asterisk. Conversion of the target base to a different nucleotide (e.g., by a point mutation) abolishes recognition/cleavage by the restriction enzymes.

In the OddCAPS method described here (Fig. 1B), three primers are designed and used together in a single PCR reaction: a “dCAPS” primer adjacent to the target base, an “Intermediate” primer also flanking the site, and a “Reverse” primer oriented opposite the others at a distance of ∼100 bp from the target. The Intermediate primer contains one or two deliberate mismatches near the target nucleotide; the dCAPS primer adds an additional one to two mismatches so that the five nucleotides at its 3’ end together with the target nucleotide form a restriction-enzyme recognition site. For amplification, the Intermediate primer is used at 1/10–1/100 of the concentration of the other primers, whereas the dCAPS and Reverse primers are supplied at standard concentrations. After PCR, the restriction enzyme is added directly to the reaction mix and the digestion products are analyzed by electrophoresis using a single gel lane (Fig. 1B).

Figure 2A shows an example of this approach applied to the *pptr-1* locus in *C. elegans*, discriminating the wild-type allele from the *pptr-1(gk257890)* allele. We introduced three nucleotide substitutions (shown in red) via the Intermediate and dCAPS primers so that only the wild-type-derived PCR product would be susceptible to NdeI cleavage. PCRs using the three primers shown in Fig. 2A were performed on genomic DNA from wild-type, *pptr-1(gk257890)* homozygotes, and wild-type/*pptr-1(gk257890)* heterozygotes; the products were digested with NdeI and separated on an agarose gel. A one-step PCR in which the Intermediate primer was used at 1/10 of the other primers’ concentration allowed unambiguous discrimination of all three genotypes: wild-type DNA yielded the NdeI-digested band pattern, *pptr-1(gk257890)* homozygous DNA remained uncut, and heterozygous DNA produced a mixture of cut and uncut bands (Fig. 2B). When the Intermediate primer was omitted, no specific PCR product was obtained (Fig. 2C), presumably because the dCAPS primer alone had too many mismatches to prime effectively. When the Intermediate primer was added at the same concentration as the dCAPS primer, even products amplified from the wild type were not cleaved (Fig. 2D), probably because preferential priming from the Intermediate primer prevented introduction of the NdeI site (Fig. 2D). To optimize the Intermediate primer concentration, we varied its relative concentration from 10^−4^ to 1; effective amplification and subsequent NdeI cleavage were observed when the Intermediate primer was present at 10^−2^ to 10^−1^ relative concentration (Fig. 2E).

**Figure 2.**
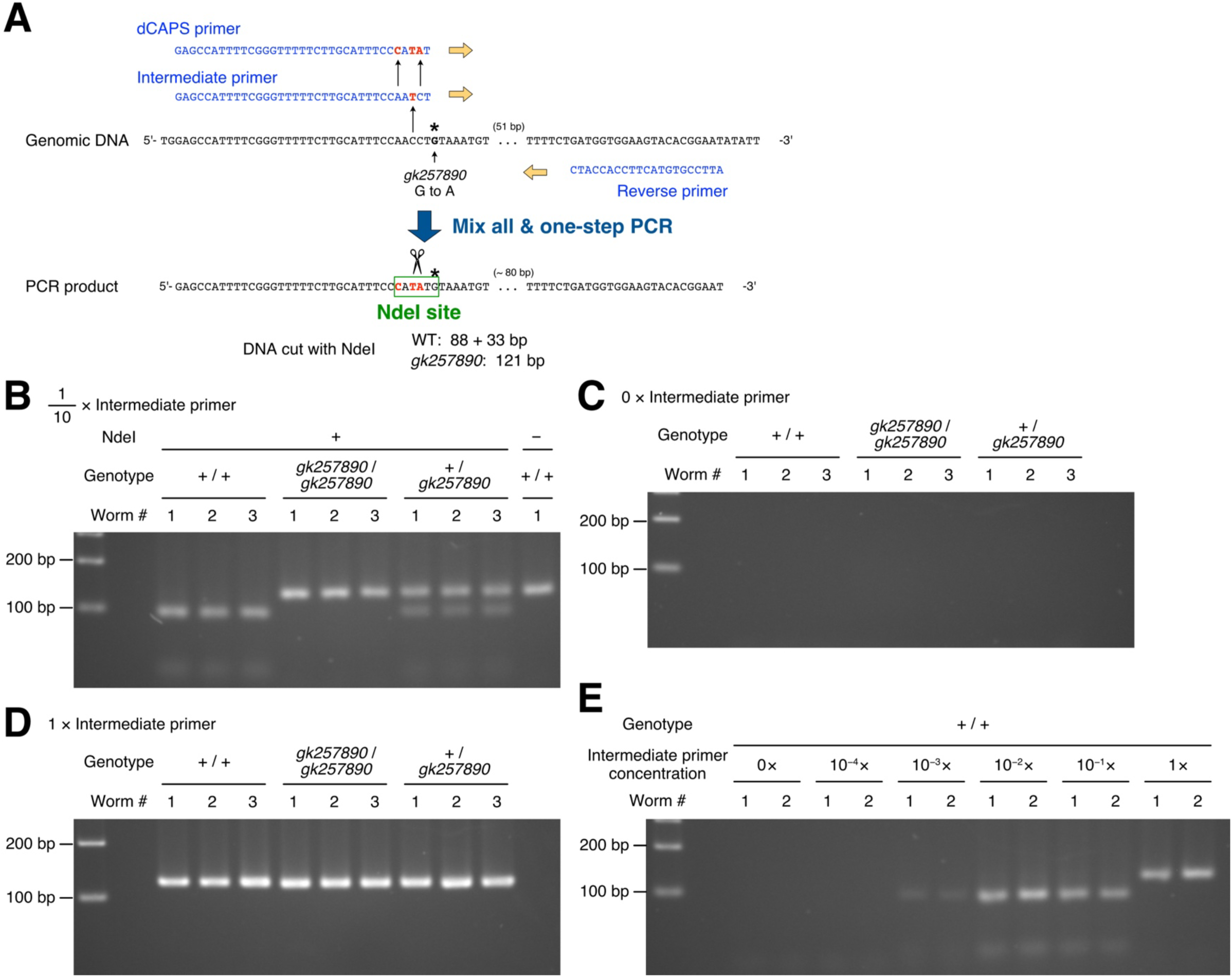
Dilution of the Intermediate primer by 10–100-fold enables one-step PCR genotyping. (**A**) Example of OddCAPS used to genotype the *C. elegans pptr-1(gk257890)* allele. In *pptr– 1(gk257890)* the G indicated by an asterisk is substituted by A. To distinguish wild-type from *pptr– 1(gk257890)*, one-step PCR was performed using an Intermediate primer that bears a single nucleotide substitution relative to the genomic sequence and a dCAPS primer that bears two additional nucleotide substitutions. The PCR product derived from wild-type DNA is cleaved by NdeI, whereas the PCR product from *pptr-1(gk257890)* is not cleaved because the allele’s substitution abolishes the NdeI recognition site. (**B**) Three individuals each of wild-type (+/+), homozygous mutant (*gk257890/gk257890*) and heterozygote (+/*gk257890*) were lysed, subjected to PCR, digested with NdeI and resolved on a 3.5% MetaPhor™ agarose gel in TAE. The Intermediate primer was included at one-tenth the concentration of the other primers. Leftmost lane, 100 bp ladder; rightmost lane, undigested wild-type PCR product. (**C**) Result when the Intermediate primer was omitted from the PCR. (**D**) Result when the Intermediate primer was included at the same (undiluted) concentration as the dCAPS and reverse primers. (**E**) The Intermediate primer concentration relative to the other primers was titrated to 0, 10^−4^, 10^−3^, 10^−2^, 10^−1^, and 1×; for each condition two animals were analyzed by PCR, NdeI digestion, and electrophoresis.

In Fig. 3, to discriminate the wild-type allele from the *dock-11(gk126522)* allele, we introduced four nucleotide substitutions stepwise into the Intermediate and dCAPS primers so that these substitutions would be incorporated into the resulting PCR product (Fig. 3A). To preserve primer extension efficiency, mismatches were not introduced at the 3′ terminal nucleotide; instead, mismatches were introduced at the fifth through second nucleotides from the 3′ end, thereby creating an EcoRI recognition site in the PCR product (Fig. 3A). Using PCR in which the Intermediate primer was present at 10% of the other primers’ concentration, followed by EcoRI digestion, we were able to identify wild-type and *dock-11(gk126522)* homozygotes as well as heterozygotes (Fig. 3B). Even with the restriction against 3′-terminal mismatches, the ability to introduce four nucleotide changes substantially expands the range of usable restriction enzymes. Theoretically, substituting the four nucleotides from the fifth to the second position from the 3′ end enables detection of any single nucleotide in a DNA molecule using a set of eight enzymes (BamHI, EcoRI, NheI, SalI, BglII, ClaI, HindIII, and MluI) (Fig. 4). The recognition sites for these eight enzymes are frequently engineered into the multiple cloning sites (MCSs) of many standard vectors; since many molecular biology laboratories are likely to already possess them, it is anticipated that experimental costs can be reduced without the need to purchase new enzymes.

**Figure 3.**
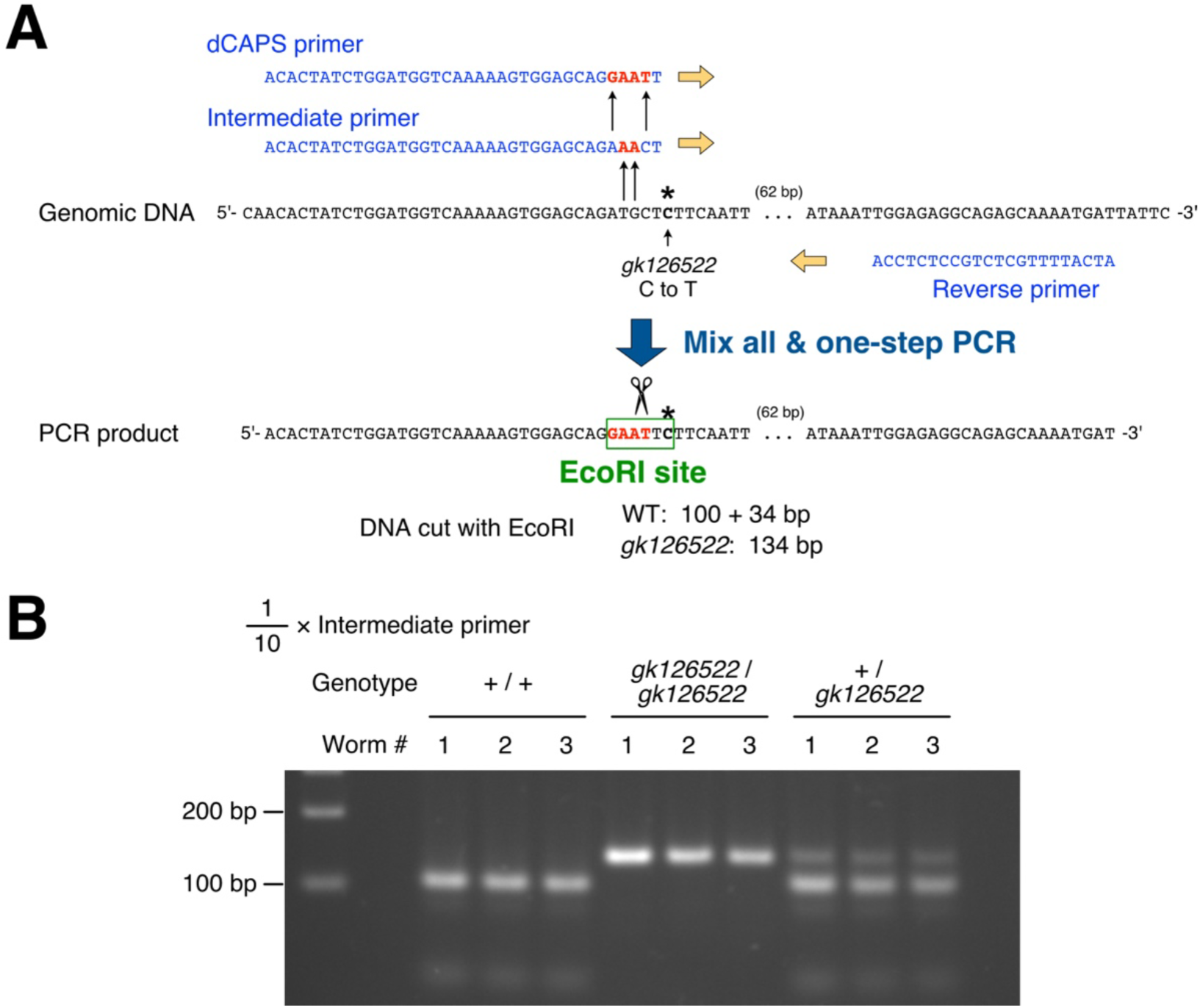
Introduction of four base substitutions by OddCAPS PCR. (A) Example of OddCAPS used to genotype the *dock-11(gk126522)* allele. In *dock-11(gk126522)* the C indicated by an asterisk is substituted by T. One-step PCR was performed using an Intermediate primer that bears two nucleotide substitutions relative to the genomic sequence and a dCAPS primer that bears two additional nucleotide substitutions. The PCR product derived from wild-type DNA is cleaved by EcoRI, whereas the PCR product from *dock-11(gk126522)* is not cleaved. (**B**) Three individuals each of wild-type (+/+), homozygous mutant (*gk126522/gk126522*) and heterozygote (+/*gk126522*) were lysed, subjected to PCR, digested with EcoRI and resolved on a 3.5% MetaPhor™ agarose gel in TAE. The Intermediate primer was included at one-tenth the concentration of the other primers. Leftmost lane, 100 bp ladder.

**Figure 4.**
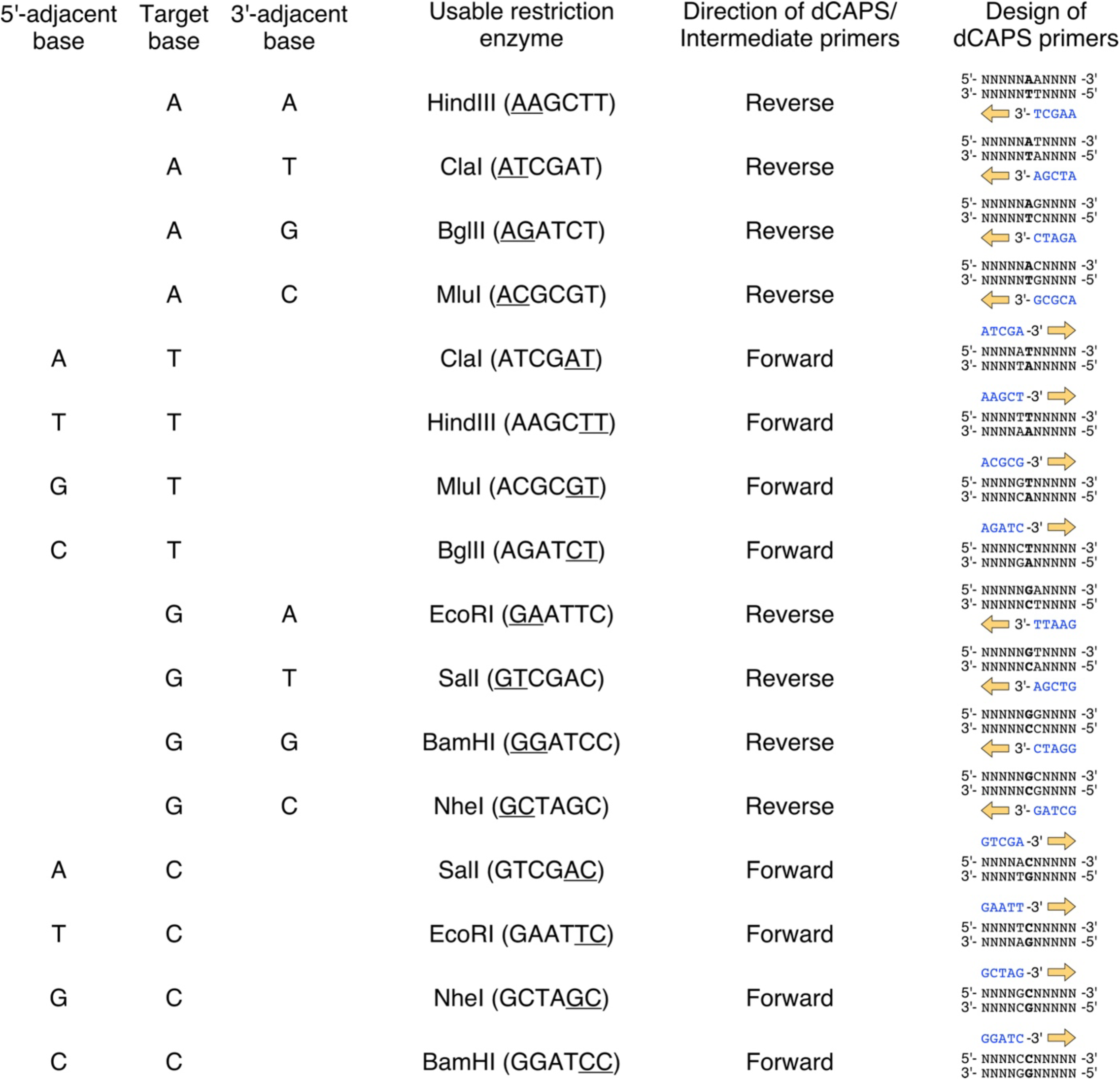
Restriction enzymes suitable for OddCAPS PCR when introducing substitutions into the four nucleotides located from five to two positions away from the target base. Using only eight commonly used restriction enzymes (BglII, ClaI, HindIII, MluI, EcoRI, BamHI, SalI, and NheI) makes it possible to detect any single nucleotide in DNA. Design the dCAPS and intermediate primers in the forward direction when the target base is T or C, and in the reverse direction when it is A or G. The sequences shown in parentheses after enzyme names are their recognition sites. Blue characters denote the five nucleotides at the 3′ end of the dCAPS primer. *Note*: We selected these eight enzymes because their cleavage sites occur in the multiple-cloning sites (MCSs) of many standard vectors, a feature that reflects their widespread use. For example, pBluescript II SK(+) includes the ClaI, HindIII, EcoRI, BamHI, and SalI sites; pEGFP N1 includes the BglII, HindIII, EcoRI, BamHI, SalI, and NheI sites; pSP73 includes the BglII, ClaI, HindIII, EcoRI, BamHI, and SalI sites; and pET 28a(+) includes the HindIII, EcoRI, BamHI, SalI, and NheI sites. Sites for MluI are also present in the MCSs of vectors such as pGL3, pCI, and pTRE. These enzymes represent common choices but are not exclusive; for example, BglII (AGATCT) may be replaced by StuI (AGGCCT), ClaI (ATCGAT) by EcoT22I (ATGCAT), MluI (ACGCGT) by SpeI (ACTAGT), EcoRI (GAATTC) by EcoRV (GATATC), BamHI (GGATCC) by KpnI (GGTACC), or NheI (GCTAGC) by SphI (GCATGC).

We also designed primers for genotyping seven additional mutant alleles—*pptr-1(dog3)*, *F36G9.13(dog4)*, *sgk-1(ft15)*, *plc-1(pe1237)*, *sulp-7(gk293386)*, *pptr-1(gk499134)*, and *him– 5(e1490)*—using OddCAPS (Fig. 5) and evaluated their effectiveness. For all these mutant alleles, OddCAPS successfully distinguished between wild-type, homozygous mutant, and heterozygous mutant genotypes (Fig. 5).

**Figure 5.**
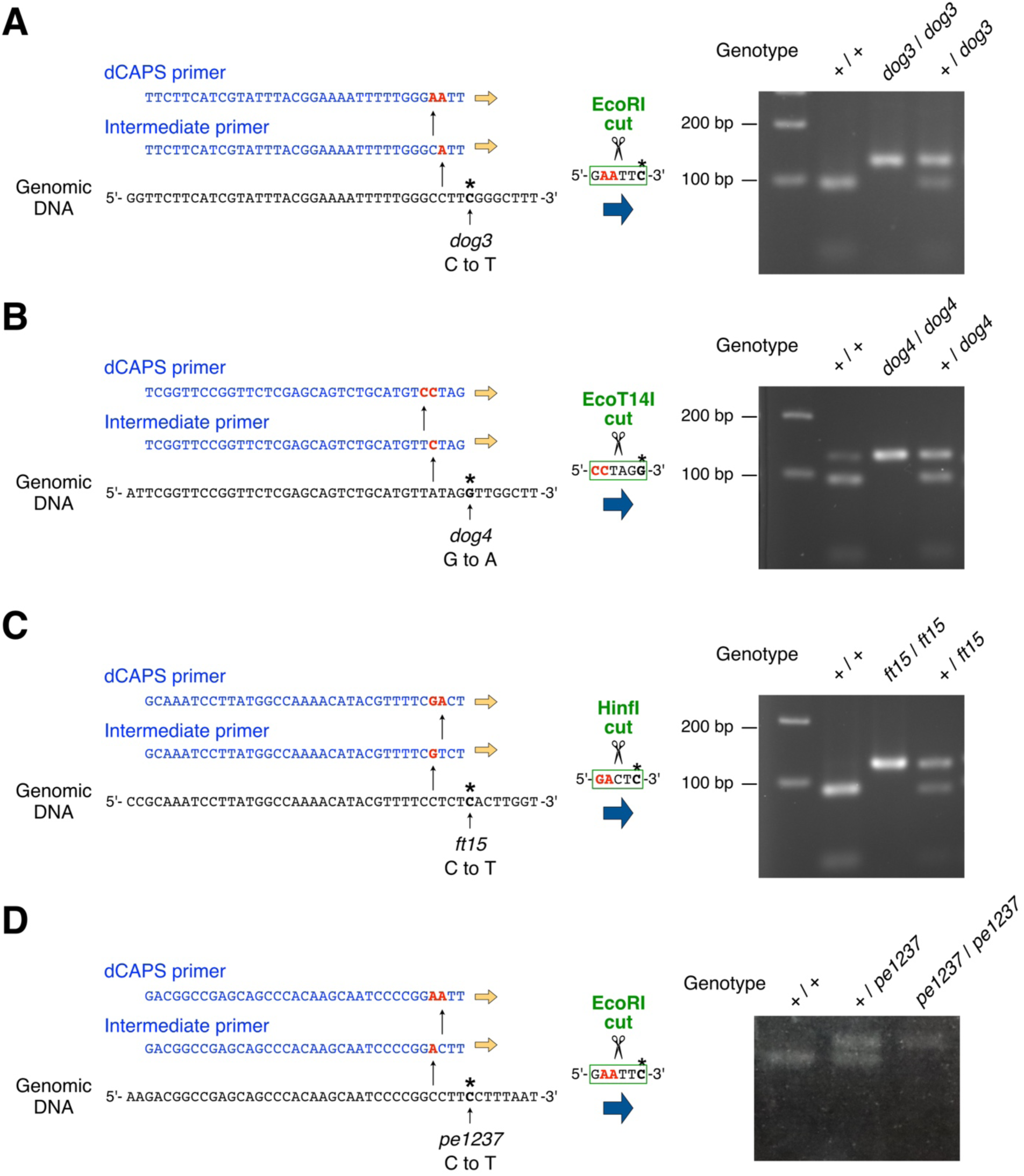
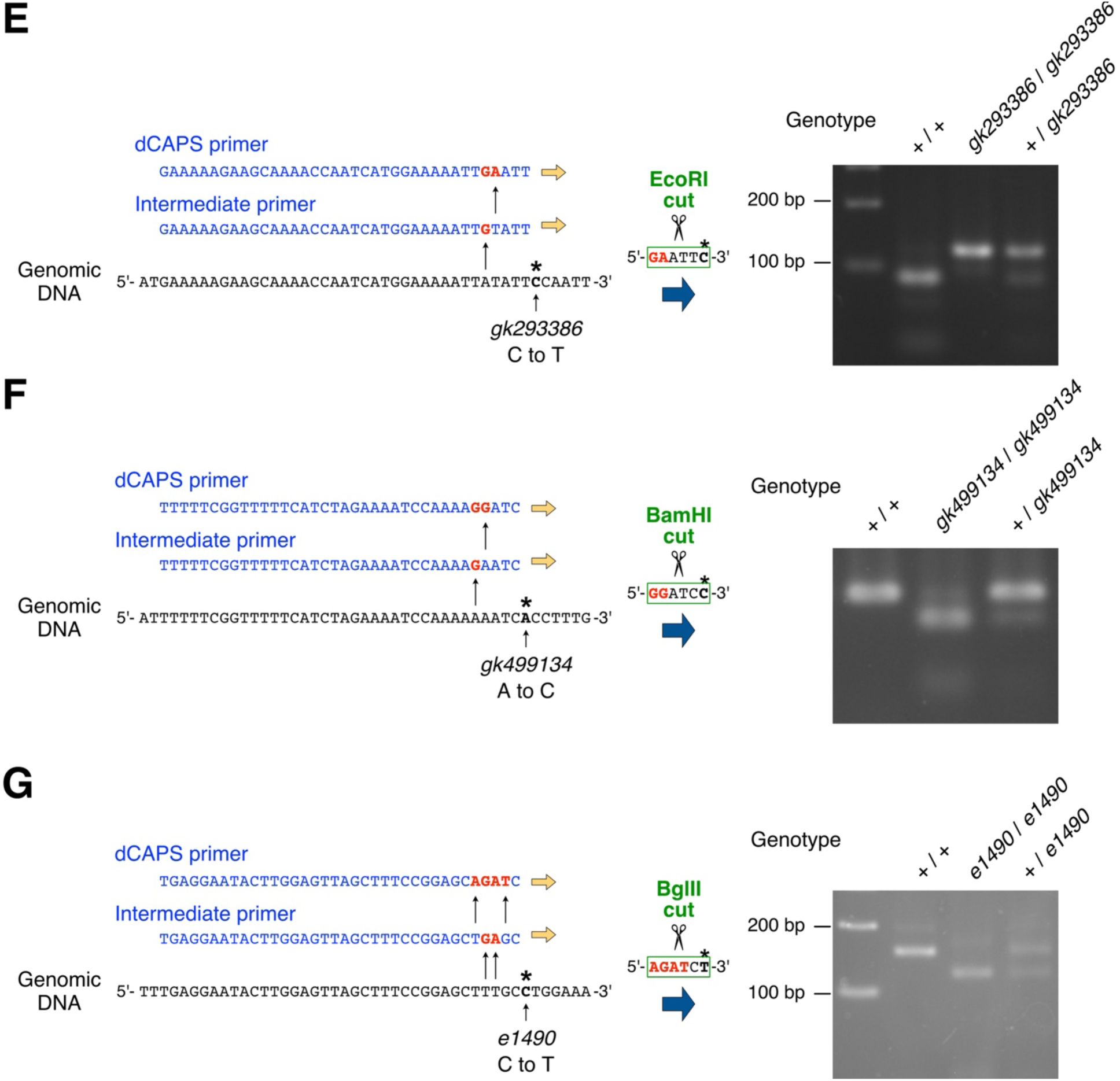
OddCAPS-based discrimination of mutant alleles. (**A**–**G**) *pptr-1(dog3)* (**A**), *F36G9.13(dog4)* (**B**), *sgk-1(ft15)* (**C**), *plc-1(pe1237)* (**D**), *sulp-7(gk293386)* (**E**), *pptr-1(gk499134)* (**F**), and *him– 5(e1490)* (**G**) were genotyped by the OddCAPS method. For each allele, OddCAPS discriminated wild-type, mutant homozygote, and mutant heterozygote samples. Panels B and F show faint residual bands likely resulting from incomplete restriction enzyme digestion; these bands partially obscured the banding patterns, but genotype discrimination remained possible for all alleles. EcoT14I and HinfI are enzymes not included in the list presented in Figure 4; they were used because they were not expensive and available in our laboratory.

The OddCAPS method enables the use of common restriction enzymes and thereby reduces the overall cost of genotyping. As an example, we evaluated the expense of genotyping 1,000 samples in our laboratory using dCAPS (Fig. 1A) versus OddCAPS (Fig. 1B). In Fig. 1A (dCAPS), primer synthesis (2 primers, total 58 nt) cost JPY 754 and MboI cost JPY 35,200 (NEB MboI [R0147S], used at 2 units per sample), giving a total of JPY 35,954 (≈ USD 225 / EUR 195 / CNY 1,514). In Fig. 1B (OddCAPS), primer synthesis (3 primers, total 94 nt) cost JPY 1,222 and EcoRI cost JPY 2,200 (NEB EcoRI-HF [R0101S], used at 2 units per sample), for a total of JPY 3,422 (≈ USD 21 / EUR 19 / CNY 144). Thus, in our setting OddCAPS reduced reagent costs by JPY 32,532 (≈ USD 204 / EUR 176 / CNY 1,370). It should be noted, however, that reagent availability and prices vary substantially between countries and laboratories, so a universally applicable and precise cost estimate is difficult to provide.

## Discussion

More than 40 years after the invention of PCR, one might ask why a strategy such as OddCAPS has not been described previously. One possible, pragmatic explanation is that changing cost structures and laboratory practices may be contributing factors. The price of custom oligonucleotide synthesis has fallen substantially, while the costs of restriction and PCR enzymes have increased. For example, EcoRI (10,000 U) listed by a Japanese supplier was JPY 6,500 in 2010 and JPY 14,000 in 2025; by contrast, another company providing oligonucleotide synthesis reported a decline from JPY 40/nt to JPY 13/nt over the same period. This creates a situation in which adding another synthesized primer is often the more economical choice than purchasing additional restriction enzymes or increasing the number of enzymatic reactions. At the same time, the adoption of high-resolution agaroses that resolve short DNAs (∼100 bp) has increased, and even simple TAE-buffered agarose electrophoresis is now capable of detecting length differences originating from primers. Finally, the rise of restriction enzyme–independent cloning methods such as Gibson assembly means that many laboratories no longer maintain large repertoires of restriction enzymes, increasing the relative advantage of the method presented here compared with dCAPS, which often depends on the use of uncommon restriction enzymes.

Another advantage of this method is that it allows straightforward transition to two-step PCR as a backup approach when the target PCR product fails to amplify by one-step PCR — although we have not observed such failures in our experience. First, perform a primary PCR using the Intermediate primer and the Reverse primer, omitting the dCAPS primer. Then use that reaction mixture directly as the template for a secondary PCR with the dCAPS primer and the Reverse primer, omitting the Intermediate primer. No additional primers need to be purchased. Because the target is short (∼100–150 bp), the likelihood of amplification failure with this two-step approach is low, although performing two PCRs requires additional time and labor. In genotyping of *pptr– 1(gk257890)* (Fig. 2), we confirmed that the two-step PCR approach also yields amplification (data not shown).

The concentration of the Intermediate primer that facilitates PCR amplification must be balanced: if it is too low, amplification can fail, whereas if it is too high, restriction enzyme cleavage of the PCR product is inhibited (Fig. 2E). However, in our genotype analysis of the *pptr– 1(gk257890)* allele, using the Intermediate primer at one-tenth the concentration of the other primers did not produce a detectable fraction of amplified wild-type DNA that remained uncut (for example, on the order of one-tenth); rather, nearly all product from wild-type DNA appeared to be cleaved (Fig. 2B, 2E). We speculate that the following sequence of events occurs during PCR. During the early PCR cycles, the Intermediate primer—having fewer mismatches with the genomic template—primarily directs a small amount of amplification. As the reaction proceeds, products bearing the substitutions introduced by the dCAPS primer become predominant, and because the dCAPS primer then outcompetes the Intermediate primer in both molecule number and annealing efficiency, amplification is progressively dominated by the dCAPS primer. As a result, the final amplicon population becomes overwhelmingly derived from the dCAPS primer.

The conventional dCAPS approach is constrained by the limited number of genomic DNA sites into which recognition sequences for commonly used restriction enzymes can be introduced; OddCAPS overcomes this limitation and allows the routine application of commonly used restriction enzymes with minimal sequence restrictions (Fig. 4). Naturally, the targeted locus must still fall within PCR-friendly parameters for GC content and repetitive elements, but because the amplicons in this method are short (∼100–150 bp), successful amplification is likely for most loci that are amenable to PCR. One practical drawback shared with dCAPS is the need for gels capable of resolving low-molecular-weight DNA, which means higher-percentage agarose or specialty products (for instance, Lonza MetaPhor™ agarose currently retails at USD 550 per 25 g). However, OddCAPS can discriminate wild-type, mutant, and heterozygous alleles in a single lane, effectively halving the number of gels required compared with many allele-specific PCR workflows. Additional cost-control strategies—using thinner gels, increasing the number of comb teeth, reusing agarose gels after melting and recasting, switching from TAE to Tris-borate-EDTA (TBE) buffer to permit lower agarose percentages, or using polyacrylamide gels—can be employed, if desired, to further reduce agarose-related costs.

*Note*: A prototype version of “OddCAPS finder,” a web browser-based program designed to assist in primer design for OddCAPS, is available at the following page (Sakai et al., manuscript in preparation): https://mcm-www.jwu.ac.jp/∼onoh/OddCAPS.html

## Materials and Methods

### Strains and culture

*C. elegans* strains were maintained by standard methods (Brenner, 1974). Worms were cultured on nematode growth medium (NGM) plates seeded with *Escherichia coli* HB101. Bristol N2 was used as the wild-type strain. Strains used in this study are listed in Table S1. For generation of *him– 5(e1490)* heterozygotes, *him-5(e1490)* males were crossed with N2 hermaphrodites and the resulting F1 males were genotyped. For heterozygotes of the other mutant alleles, CAT117 males (*Ex*[*myo-3p::venus*]), which express green fluorescent protein in body-wall muscle, were crossed to hermaphrodites of each mutant; F1 hermaphrodites showing *Ex*[*myo-3p::venus*] fluorescence, indicative of cross progeny, were genotyped.

### Single worm genotyping

Genotyping by single-worm lysis, PCR, and restriction digestion was performed with modifications to the method of Wicks et al. (Wicks et al., 2001). A detailed protocol, from primer design to genotype detection by electrophoresis, is available in the Supplemental Text 1. In brief, one adult worm was transferred to an NGM plate seeded with HB101 and allowed to lay eggs overnight. The adult was lysed in 10 µL of single-worm lysis buffer (50 mM KCl, 10 mM Tris-HCl [pH 8.3], 2.5 mM MgCl_2_, 0.45% NP-40, 0.45% Tween 20, 0.01% gelatin, and 60 µg/mL Proteinase K), and 1 µL of this lysate served as the template for PCR using Quick Taq HS DyeMix (TOYOBO, Cat. No. DTM-101). Custom oligonucleotide primers were obtained from Integrated DNA Technologies (IDT) at the standard desalting grade and used without further purification. After PCR, 0.3 µL of the selected restriction enzyme plus the appropriate restriction buffer was added directly to the reaction and incubated overnight; digestion products were run directly on agarose gels. Electrophoresis was carried out on 3.5% MetaPhor™ agarose (Lonza, Cat. No. 50180) in TAE buffer, with an ExcelBand 100 bp DNA Ladder (SMOBIO, Cat. No. DM2100). Gels were stained in TAE containing 0.5 µg/mL ethidium bromide (Nacalai Tesque, Inc., Cat. No. 14631-94), visualized using a UV transilluminator (WUV-M20, ATTO, Tokyo, Japan) and imaged using a gel documentation system (Printgraph AE-6932, ATTO, Tokyo, Japan). Primers used in this study are listed in Table S2. Frequently asked questions and answers are available in the Supplemental Text 2.

## Acknowledgments

We thank the *Caenorhabditis* Genetics Center (funded by the NIH Office of Research Infrastructure Programs P40 OD010440) and Dr. Hirofumi Kunitomo for strains. This work was supported by JSPS KAKENHI Grant Numbers JP25KJ2254 to N.S. and JP26K09212 to H.O. The authors have no competing interests. The raw data is available on figshare: https://doi.org/10.6084/m9.figshare.33085313.

## Figures and tables

**Table S1:**
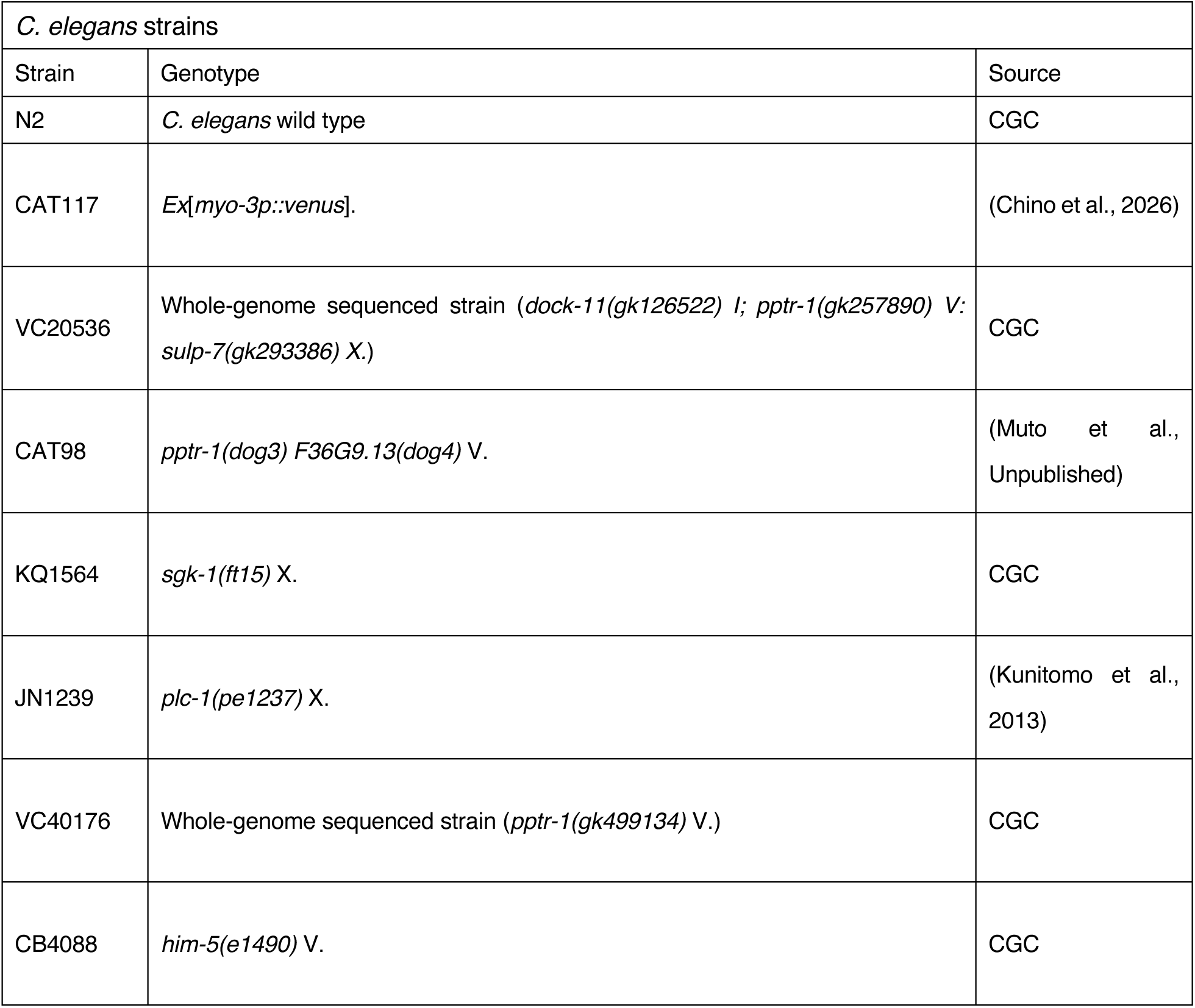
Strains used in this study.

**Table S2:**
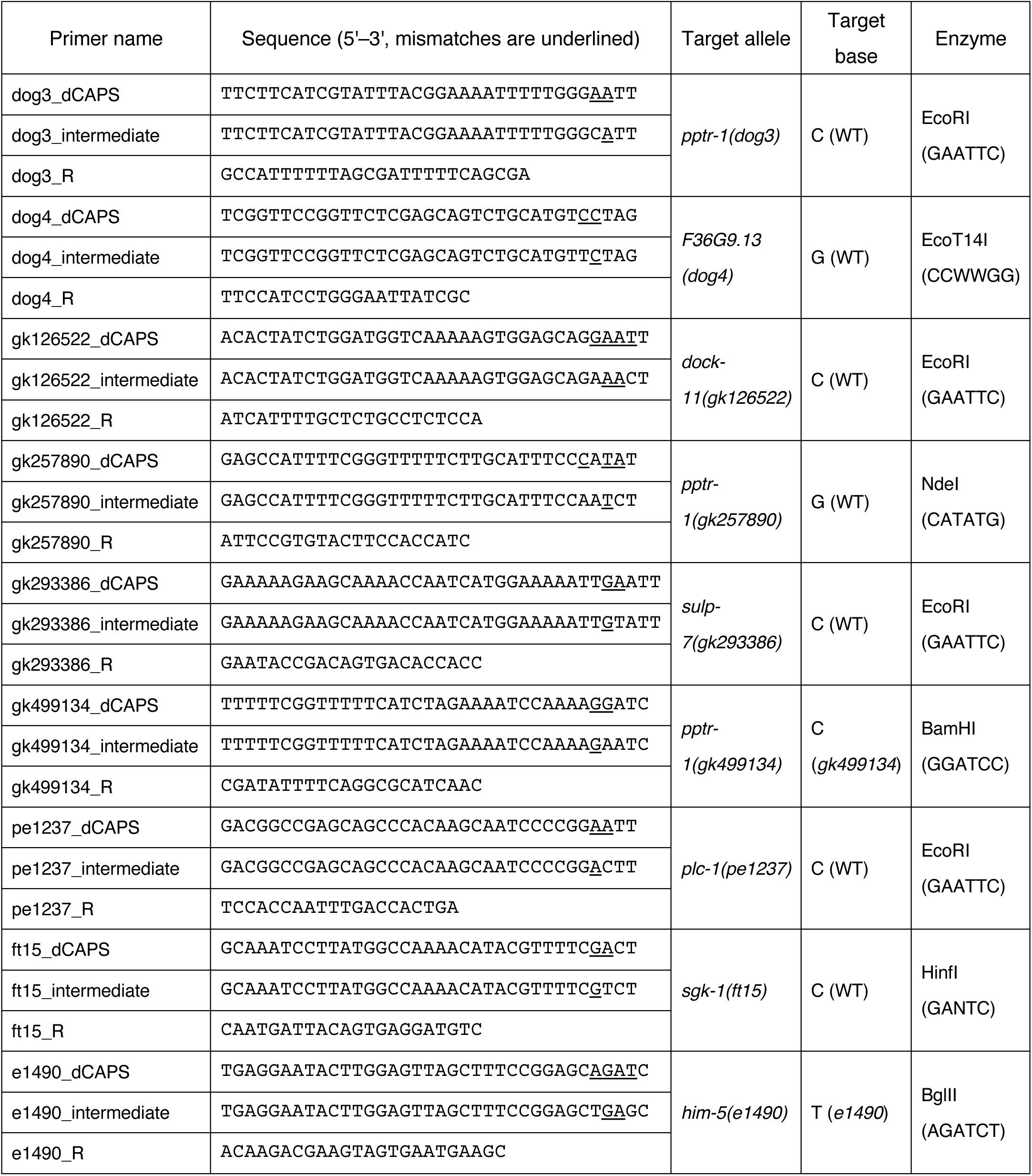
Primers used in this study.

## Supplemental Text 1

### Protocol from primer design through genotype detection by gel electrophoresis

#### A. Primer design and preparation

As an example, we sought to distinguish the wild-type *C. elegans* allele from the *pptr-1(gk257890)* mutant (Fig. 3A). The sequence on chromosome V that corresponds to positions 16,359,127– 16,359,141 (presented here in the orientation opposite to the gene) is 5’-CCAACCT**<u>G</u>**TAAATGT-3’ in the wild type. In the *pptr-1(gk257890)* allele, the central guanine (G) is substituted by adenine (A), producing 5’-CCAACCT**<u>A</u>**TAAATGT-3’ (Fig. 3A).

1. Open dCAPS Finder 2.0 (Neff et al., 2002; http://helix.wustl.edu/dcaps/). Enter the wild-type sequence “CCAACCTGTAAATGT” into the field labeled “Enter the Wild Type Sequence”, and enter the mutant sequence “CCAACCTATAAATGT” into “Enter the Mutant Sequence”. Set “How many mismatches in the primer?” to “0” and click “submit”. *Note*: The ApE program (Davis & Jorgensen, 2022; https://jorgensen.biology.utah.edu/wayned/ape/) also includes a comparable “dCAPS calculator” function.
2. If you are fortunate to find a suitable restriction enzyme that you routinely use, perform PCR-RFLP (CAPS) with that enzyme. If no satisfactory enzyme is returned, go back to the first dCAPS Finder page, increase the value of “How many mismatches in the primer?” to “1”, and press “submit”.
3. If you are fortunate enough to identify a satisfactory restriction enzyme and the corresponding “PRIMER SEQUENCE” does not contain a mismatch at the 3′-terminal nucleotide (underlined in the output), perform the dCAPS assay using that enzyme and primer. If no suitable enzyme is found, return to the initial screen, set “How many mismatches in the primer?” to “2”, and click “submit”. *Note*: The 3′-terminal nucleotide is important for efficient PCR amplification; it is advisable to avoid introducing mismatches at that position.
4. Continue increasing the number of primer mismatches one increment at a time until a satisfactory restriction enzyme is identified. If the number of mismatches reaches two or greater, proceed using the OddCAPS approach described below. If permitting three mismatches still fails to identify an effective primer/enzyme pair, select one of the restriction enzymes listed in Figure 4 and design a dCAPS primer that incorporates the four specific mismatches indicated for that enzyme (see Figure 4). *Note*: As mentioned in the main text, a prototype version of “OddCAPS finder,” a web browser-based program designed to assist in primer design for OddCAPS, is available at the following page (Sakai et al., manuscript in preparation): https://mcm-www.jwu.ac.jp/∼onoh/OddCAPS.html
5. When two or more mismatches are necessary, implement the OddCAPS approach. Design a dCAPS primer of approximately 35 nucleotides in length that introduces 2–4 mismatches relative to the genomic template within the region spanning the fifth to the second nucleotide from the 3′ end, so that PCR amplification will create a restriction-enzyme recognition site (Figure 2A). The primer’s terminal five nucleotides (for a 6-base cutter), together with the targeted base, should constitute the restriction-enzyme recognition sequence; when the target base is substituted, the site will be absent and the PCR product will therefore be resistant to digestion.
6. Design an Intermediate primer that serves as a stepping-stone between the genomic sequence and the dCAPS primer. Design the Intermediate primer so that it differs from the genomic DNA by 1–2 nucleotides, and differs from the dCAPS primer by an additional 1–2 nucleotides, thereby increasing mismatches incrementally. For the intermediate primer, avoid introducing mismatches in the two terminal nucleotides at the 3′ end, since these bases may be important for efficient PCR initiation. Place the mismatch at the second nucleotide from the 3′ end in the dCAPS primer rather than in the intermediate primer (Figs. 2A and 3A). *Note*: Although untested, an intermediate primer with a slightly truncated 5′ end is unlikely to prevent PCR amplification and subsequent genotyping. In contrast, a longer dCAPS primer can improve the visual distinction between cut and uncut fragments following restriction digestion; this can allow the use of lower-concentration agarose gels, potentially resulting in further cost savings.
7. Design a Reverse primer in the antisense orientation located approximately 80–150 bp downstream of the dCAPS primer. Unlike the dCAPS or Intermediate primers, the reverse primer’s exact placement is flexible; aim for a primer 20–25 nt in length with a GC content near 40–60%. To maximize the likelihood of successful PCR, employ primer-design tools (for example, Primer-BLAST at https://www.ncbi.nlm.nih.gov/tools/primer-blast/index.cgi).
8. Confirm that the restriction site for your selected enzyme is not located in the vicinity of the target base. A proximal recognition sequence can cause the PCR amplicon to be cut irrespective of the allelic state, preventing reliable genotyping. If such a nearby site is present, choose a different restriction enzyme.
9. Synthesize the dCAPS, Intermediate, and Reverse primers through a commercial oligonucleotide synthesis service (e.g., Integrated DNA Technologies). Resuspend each lyophilized oligo to a final concentration of 100 µM in water or TE buffer.

#### B. Worm lysis

*Note*: The following worm-lysis procedure is a modification of the method of Wicks et al. (Wicks et al., 2001) for genotyping *Caenorhabditis elegans*. For other organisms, standard genotyping methods (for example, mouse tail biopsy) may be substituted.

1. Prepare a single-worm lysis buffer composed of 50 mM KCl, 10 mM Tris-HCl (pH 8.3), 2.5 mM MgCl_2_, 0.45% NP-40, 0.45% Tween 20, and 0.01% gelatine. Sterilize the buffer by autoclaving.
2. Just before use, add 3 µL of 20 mg/mL proteinase K to 1 mL of single-worm lysis buffer to achieve a final concentration of 60 µg/mL.
3. Dispense 10 µL of single-worm lysis buffer (+ proteinase K) into each well of a 96-well PCR plate or into individual PCR tubes.
4. Using a stereomicroscope, transfer one worm to be genotyped (if you plan to use the progeny after genotyping, transfer a parent worm isolated on a plate and allowed to lay eggs overnight) into each well, taking care not to touch the sidewall of the well when depositing the worm.
5. Seal the plate carefully with PCR-grade adhesive film to prevent evaporation.
6. Lyse the worms in a thermocycler using the following program: 60°C for 60 min; 95°C for 15 min; hold at 15°C. The crude DNA lysate can be stored at 4°C for several days or at −20°C for several months.

#### C. PCR

1. While gently mixing by pipetting, transfer 1 µL of each crude DNA lysate from section B into individual wells of a separate 96-well PCR plate.
2. Add 10 µL of the following PCR mix to each well. PCR mix (per sample, total: 10 µL)
  - H_2_O: 4.85 µL
  - 2× Quick Taq HS DyeMix (TOYOBO, Cat. No. DTM-101): 5 µL
  - 100 µM dCAPS primer: 0.05 µL
  - 10 µM (or 1 µM) Intermediate primer: 0.05 µL
  - 100 µM Reverse primer: 0.05 µL *Note*: An alternative product comparable to Quick Taq is GoTaq® Green Master Mix (Promega, Cat. No. M7123).
3. Seal the plate carefully with PCR-grade adhesive film to prevent evaporation.
4. Run the following thermal-cycler program: 94°C for 2 min; 40 cycles of 94°C for 30 sec, 58°C for 30 sec, 68°C for 20 sec; 68°C for 2 min; 15°C hold. *Note*: We have not experimentally verified the benefit, but we employ a slightly elevated cycle number to offset a potential slow PCR ramp-up. *Note*: The post-PCR reaction solution is stable at 4°C for several days and can be stored at −20°C for extended periods.

#### D. Restriction enzyme digestion

1. Add 10 µL of the following restriction cocktail to each PCR well. Restriction cocktail (per sample, total 10 µL)
  - H_2_O: 7.7 µL
  - 10× restriction enzyme buffer (appropriate for the enzyme): 2.0 µL
  - Restriction enzyme: 0.3 µL
2. Seal the plate with a fresh PCR adhesive film to prevent evaporation, briefly tap to mix, and incubate at the enzyme’s recommended reaction temperature for several hours to overnight.

#### E. Electrophoresis

- Apply the restriction enzyme reaction mixtures directly to a 3.5% MetaPhor™ agarose gel cast in TAE buffer and resolve the fragments by standard gel electrophoresis. When running large numbers of samples, consider using a high-throughput gel platform that accepts direct loading from 8– or 12-channel pipettes (e.g., Nihon Eido NB-1017B).
- Stain the gel in TAE buffer containing 0.5 µg/mL ethidium bromide and visualize bands using a UV transilluminator. *Note*: We have not tried it, but the OddCAPS method should also be applicable to short insertions and deletions (indels).

## Supplemental Text 2

### Frequently Asked Questions

1. Can costs be reduced further? Possible cost-reduction strategies include: substituting TBE for TAE to allow use of lower-percentage agarose gels; casting thinner gels to decrease agarose consumption; adopting multi-sample high-throughput electrophoresis platforms; shortening the Intermediate primer to reduce oligonucleotide synthesis expense; lengthening dCAPS primers at the 5’ end to permit use of lower-concentration agarose gels; and preparing Taq polymerase in-house rather than purchasing commercial enzyme. As an optional, modest additional step, one can also heat the used agarose gel so that it melts again, pass it through a paper filter (for example, kitchen oil-filter paper), and then allow it to solidify again for reuse (http://rizo-inc.cocolog-nifty.com/blog/2011/09/post-85ef.html).
2. Which allele should be designed to be cleavable, wild-type or mutant? It depends on your objective. In general, it is better to design the assay so that the allele whose false-positive identification would be most problematic is the one that is cleaved. This is because failure to observe cleavage can result from technical error (for example, forgetting to add the restriction enzyme). For example, if your aim is to isolate mutant homozygotes after a cross, it is better to introduce a restriction site only in the mutant DNA to minimize false positives. Conversely, if you wish to select wild-type homozygotes (i.e., remove the mutation), it is better to produce a site that is cleaved in the wild-type DNA.
3. How can one resolve DNA fragments of roughly 100 bp by electrophoresis? We have obtained reliable separation using high-percentage agarose gels, for example 3.5% MetaPhor™ agarose (Lonza, Cat. No. 50180) or 5% NuSieve™ 3:1 agarose (Lonza, Cat. No. 50090). Polyacrylamide gel electrophoresis is also a suitable alternative.
4. Which DNA polymerase should be used for PCR? We recommend using a conventional, inexpensive Taq DNA polymerase rather than a high-performance, proofreading enzyme. Standard Taq is cost-effective, its reaction buffer is less likely to contain components that inhibit downstream restriction digests, and its lack of 3’→5’ exonuclease (proofreading) activity reduces the chance that primer-introduced mismatches will be corrected during amplification.
5. Which Intermediate primer concentration is preferable, 1/10 or 1/100? We use a 1:10 dilution of the Intermediate primer when the design introduces a total of 3–4 mismatches; the higher primer concentration helps ensure robust amplification under these more challenging conditions. When only 1–2 mismatches are introduced, reduce the Intermediate primer to a 1:100 dilution to promote complete restriction-enzyme digestion of the PCR product.
6. What should I do if PCR fails to yield the expected product? Although we have not observed amplification failures with OddCAPS in our laboratory, some genomic loci may be refractory to PCR. Rather than extensive empirical optimization (e.g., iteratively varying annealing temperature or primer concentration), we recommend two-step PCR (see Discussion).

